# CIPHER: An end-to-end framework for designing optimized aggregated spatial transcriptomics experiments

**DOI:** 10.64898/2026.01.08.698503

**Authors:** Zachery Hemminger, Haley De Ocampo, Fangming Xie, Zhiqian Zhai, Jingyi Jessica Li, Roy Wollman

## Abstract

**Motivation:** Most imaging-based spatial transcriptomics methods measure individual genes, which limits scalability and typically requires integration with scRNA-seq to recover full cellular states. Recent approaches such as CISI, FISHnCHIPs, and ATLAS address this limitation by measuring aggregate transcriptional signatures, where multiple genes are pooled into each channel to increase throughput. While aggregate measurements improve scalability, they shift the problem from gene selection to feature design. For effective integration with scRNA-seq, these signatures must be not only discriminative in transcriptional space but also straightforward to measure, with balanced signal, sufficient dynamic range, and robustness to experimental noise. By optimizing decoding accuracy in isolation, existing methods leave substantial performance on the table.

**Results:** We present CIPHER (Cell Identity Projection using Hybridization Encoding Rules), a neural-network framework that jointly optimizes the experimental encoding matrix, i.e., the way that genes are aggregated to signatures, and the downstream cell embedding. CIPHER integrates the physical limits of imaging assays directly into its loss function, shaping the latent space to maximize discriminability while maintaining robustness to measurement noise and signal constraints. Using a large-scale mouse brain scRNA-seq reference, we show that CIPHER-designed encodings yield latent spaces with improved cell-type separability, uniform signal utilization, and greater resilience to hybridization variability, resulting in higher decoding accuracy from both simulated and experimental data.

**Conclusion:** CIPHER formulates aggregate signature design as a joint optimization problem over decoding accuracy and experimental measurability. This enables systematic, scRNA-seq-aligned feature design for scalable spatial transcriptomics based on aggregate measurements.

**Availability:** Code and documentation are available at https://github.com/wollmanlab/Design/.

**Author summary:** Spatial transcriptomics reveals how cells are organized within tissues by mapping where genes are expressed. To achieve both scale and resolution, many approaches now combine spatial imaging with single-cell RNA-seq references to reconstruct complete transcriptomes *in situ*. New methods such as CISI, FISHnCHIPs, and ATLAS accelerate this process by measuring combinations of co-expressed genes rather than each gene individually. These aggregate measurements simplify experiments but introduce a new challenge: deciding which genes to combine so that the resulting features are both experimentally reliable and computationally informative for integration with scRNA-seq data. We developed CIPHER, a computational framework that learns how to design and decode these aggregate measurements optimally. By integrating experimental constraints and decoding accuracy into a unified neural-network model, CIPHER provides a principled approach for designing signature-based spatial transcriptomics experiments that enable efficient and accurate reconstruction of cellular transcriptomes.

## Introduction

Spatial transcriptomics aims to connect gene expression with cellular organization in intact tissues, but existing platforms face fundamental trade-offs between throughput, sensitivity, and spatial resolution [1, 2]. Imaging-based approaches that directly detect individual RNA molecules can achieve high spatial precision but scale poorly. High-magnification, spot-based methods require long acquisition times and are limited to small fields of view [3–5], while lower-magnification imaging platforms trade molecular resolution for speed and coverage [6, 7]. Sequencing-based spatial assays avoid optical crowding altogether but capture only a small fraction of the transcriptome per spatial unit, with detection biased toward abundant transcripts [2, 8, 9]. As a result, no current technology can simultaneously achieve high sensitivity, large spatial coverage, and comprehensive transcriptome profiling. Most spatial transcriptomics studies therefore rely on integration with scRNAseq data to recover cell identities and transcriptional states, placing increasing emphasis on how spatial measurements are designed to support downstream integration rather than direct gene-by-gene readout.

To overcome the limitations of gene-by-gene spatial measurements, a new class of spatial transcriptomics methods adopts a different objective: designing measurements that enable easy integration with scRNAseq references rather than attempting to directly recover partial transcriptomes *in situ*. Methods such as CISI, FISHnCHIPs, and ATLAS replace individual gene measurements with aggregate transcriptional signatures, in which multiple genes are combined into a smaller number of composite features [10–12]. By leveraging prior knowledge of transcriptional structure learned from single-cell data, these approaches shift the burden of resolution from the imaging assay to the computational decoding step. This reframing enables higher throughput, reduced imaging demands, and more efficient use of experimental signal while preserving the ability to assign cells to reference-defined identities and states.

Although CISI, FISHnCHIPs, and ATLAS share the goal of integration-first spatial measurement, they differ fundamentally in how aggregate encodings are constructed and interpreted. CISI introduced the use of composite measurements by applying compressed sensing to randomly generated gene mixtures, followed by sparse recovery to reconstruct individual gene expression profiles [10]. FISHnCHIPs instead defines aggregate features by grouping co-expressed genes into modules derived from single-cell data, prioritizing signal amplification and sensitivity through coordinated hybridization [11]. ATLAS generalizes aggregate measurement design by representing each feature as a weighted combination of genes and explicitly learning a linear mapping from gene expression space into a low-dimensional latent space aligned with known cell types [12]. This formulation enables supervised optimization of encodings for cell-type separability and scalability, allowing ATLAS to profile tens of millions of cells across hundreds of tissue sections while maintaining alignment with single-cell references [12]. Across all three approaches, aggregate measurement design follows a common workflow of computational encoding, *in situ* measurement, and computational decoding, but differs in how the latent space is defined and optimized.

Despite their success, existing aggregate spatial transcriptomics methods do not explicitly model the experimental factors that determine whether an encoding can be measured reliably. Encoding design is typically performed entirely in silico, without accounting for finite signal dynamic range, variability in hybridization efficiency across probes, background structure, or uneven signal allocation across channels. These effects directly shape signal separability and decoding performance in real experiments, yet they are absent from the encoding optimization itself [3, 4, 11]. In practice, this leads to encodings that may be highly discriminative in theory but are difficult to read out robustly, or that waste measurement capacity on features that are poorly constrained by the assay. Addressing this gap requires design frameworks that treat physical and biochemical measurement limits as first-class components of the encoding problem.

Here we present CIPHER (Cell Identity Projection using Hybridization Encoding Rules), a computational framework for designing aggregate spatial transcriptomics encodings that explicitly account for experimental measurability. CIPHER jointly optimizes encoding and decoding accuracy under physical constraints on signal dynamic range, probe usage, and measurement robustness. We first introduce the conceptual framework underlying measurability-aware encoding design (Figure 1), then quantify the trade-offs imposed by experimental constraints and their impact on encoding performance (Figures 2–4). We compare CIPHER-designed encodings to existing strategies, including DPNMF, the method used in the ATLAS paper [12](Figure 4), and evaluate performance using large-scale simulations (Figure 6) and experimental validation in mouse brain tissue (Figure 7). Finally, we demonstrate how CIPHER can be applied to additional organs beyond the mouse brain (Figure 8). Together, these results establish CIPHER as a principled framework for information-optimal and experimentally feasible design in signature-based spatial transcriptomics.

**Fig 1.**
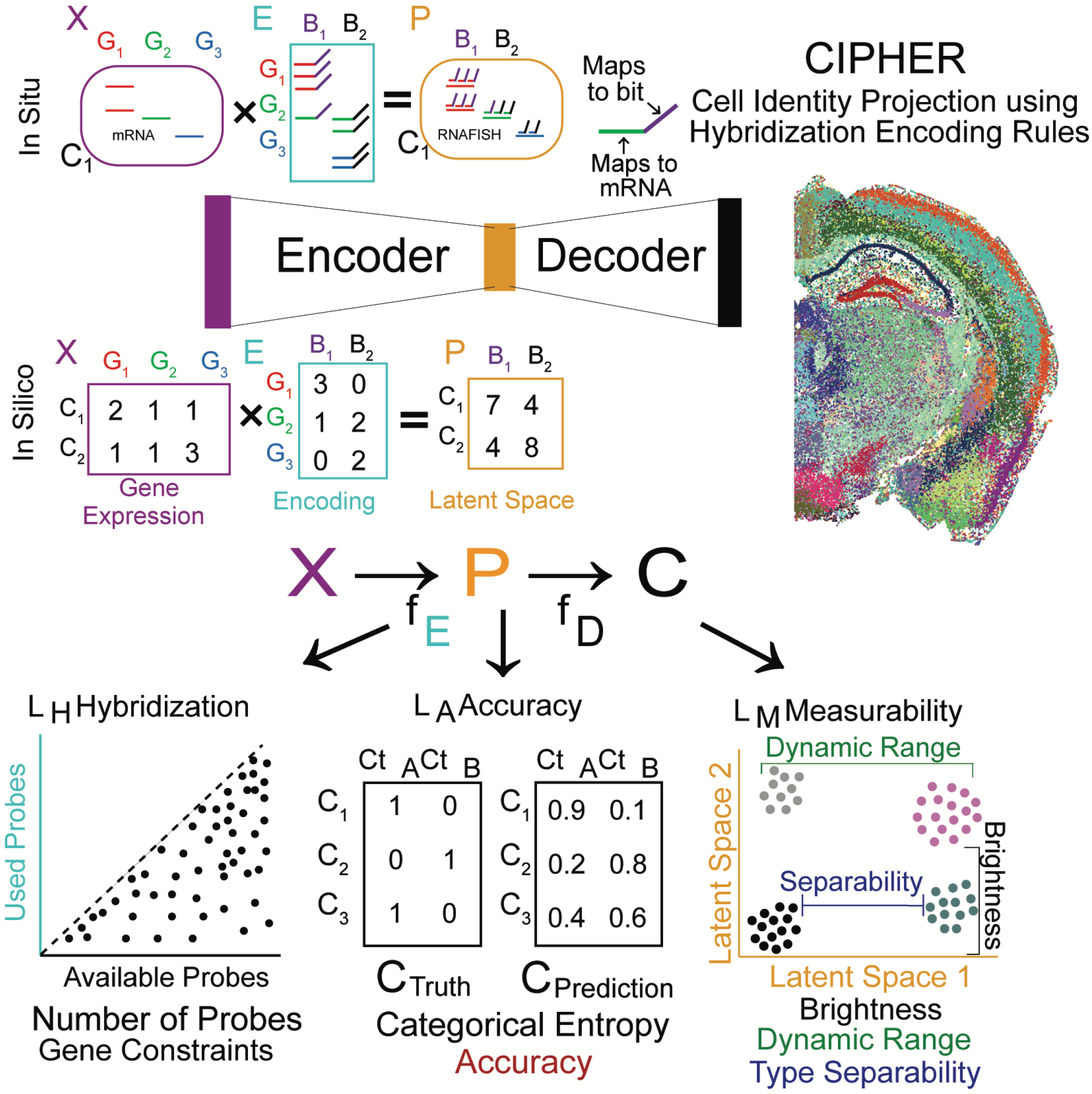
The CIPHER Framework. Schematic of the end-to-end neural-network for aggregate spatial transcriptomics design. The top panel illustrates the principle of aggregate FISH, demonstrating how molecular probes physically implement a linear weighted-sum projection of gene expression *in situ* prior to measurement. The bottom panel details the CIPHER framework, which employs a machine learning approach to fit the optimal encoder matrix (*E*) using single-cell RNA-seq reference inputs (*X*). This optimization maximizes decoding accuracy (*L*_*A*_) while strictly enforcing physical hybridization constraints (*L*_*H*_) and ensuring experimental measurability (*L*_*M*_), yielding a latent space (*P*) that is both informative and physically realizable. The decoder (*D*) subsequently reconstructs cell identity (*C*) from these optimized signals.

**Fig 2.**
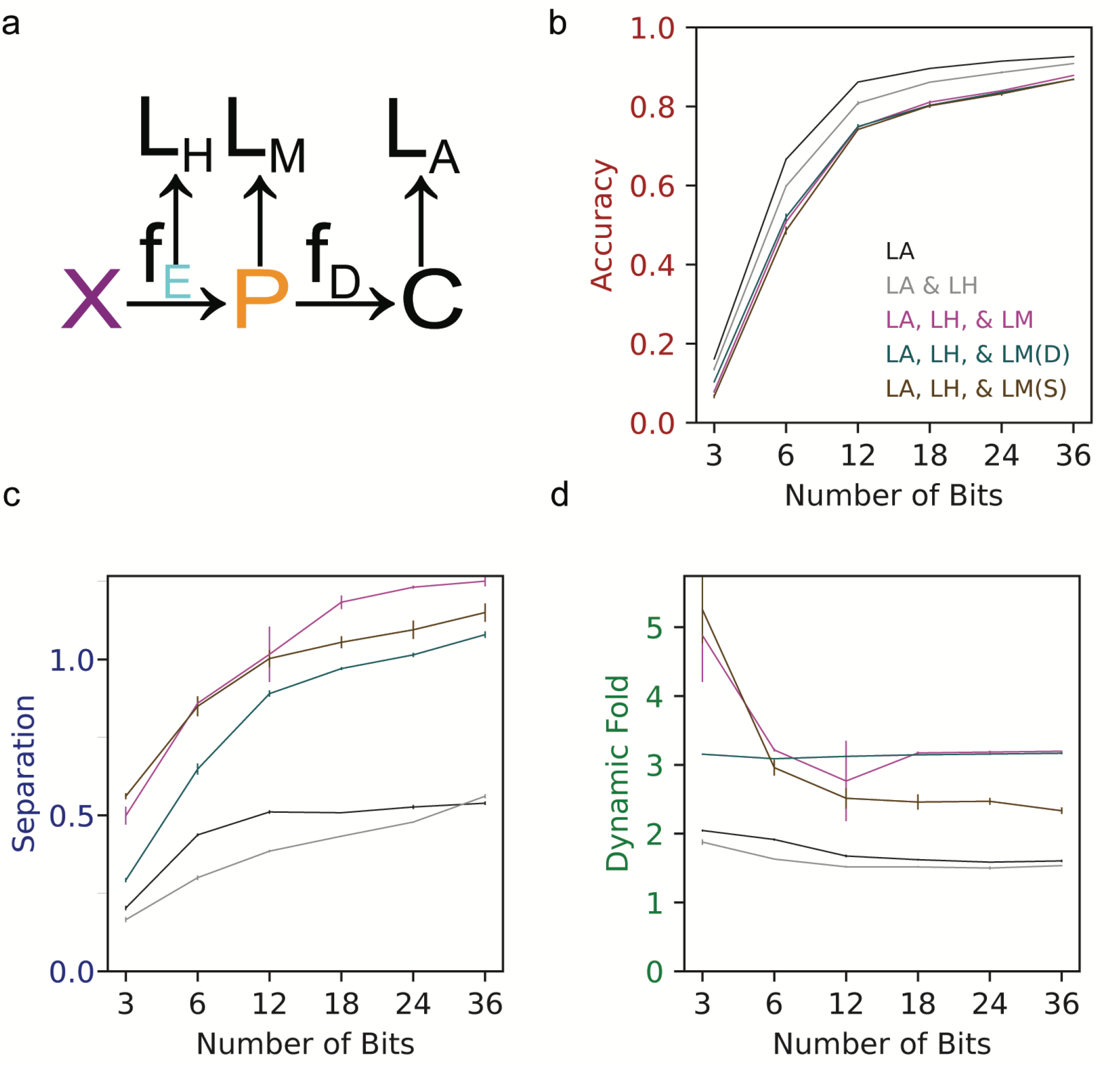
Impact of Loss Functions on Encoding Performance. (A) Schematic of the composite loss function components. *L*_*H*_ (Hybridization Rules) enforces physical probe constraints such as global probe budgets and per-gene limits. *L*_*M*_ (Measurability) enforces sufficient dynamic fold and cell-type separation, as well as overall brightness. *L*_*A*_ (Accuracy) maximizes cell-type classification performance. (B–D) Performance metrics plotted against the number of bits. All three panels display the same five loss-function configurations to isolate the specific impact of each term: Accuracy only (*L*_*A*_), Accuracy + Hybridization (*L*_*A*_, *L*_*H*_), and the full model (*L*_*A*_, *L*_*H*_, *L*_*M*_), including variants with only the Dynamic Range or Separability components of *L*_*M*_ enabled. (B) Classification accuracy shows a modest decrease with the addition of measurability terms. This trade-off yields substantial gains in (C) latent-space separability and (D) dynamic fold change, demonstrating how *L*_*M*_ enforces sufficient signal dynamic range for robust experimental readout compared to unconstrained encoders. See Methods for the definitions of Accuracy, Separation, and Dynamic Fold.

## Methods

### Terminology

We use the term *bit* to denote a single dimension of the encoded measurement space. Unlike classical MERFISH-style bits, which represent binary presence or absence of signal, CIPHER bits correspond to continuous-valued aggregate transcriptional features formed by pooling multiple genes into a single readout channel.

### Datasets and Preprocessing

CIPHER was evaluated across three biological contexts: adult mouse brain, developing mouse, and mouse lymph node. For all datasets, raw scRNA-seq count matrices were processed uniformly. Genes were retained if probe designs were available in the PaintSHOP probe database; no additional filtering based on expression level, transcript length, or sequence complexity was applied. Mitochondrial and ribosomal genes were retained if probe designs were available. Gene expression values were normalized to counts per 100,000 (CP100k) by dividing each cell by its total UMI count and multiplying by 10^5^. Since normalized values were not rounded, this differs from CP10k only by a global factor of 10 and serves mainly to place projected bit intensities and brightness targets in a convenient numerical range.

#### Adult mouse brain

We used the Allen Brain Cell Atlas whole mouse brain 10X scRNA-seq dataset (WMB-10X) [13]. Cells with valid metadata and genes shared across 10X v2 and v3 chemistries were retained. To ensure balanced representation during training, the dataset was subsampled to 100,000 cells using stratified sampling, with oversampling of rare cell types. Cell-type labels correspond to the subclass annotation level (*K* = 336 categories). The Allen Brain Atlas subclass distribution is highly imbalanced, with a small number of abundant non-neuronal subclasses comprising a disproportionately large fraction of all cells. Without stratified sampling, optimization would be dominated by these abundant populations and would underweight the many rarer neuronal subclasses that are central to the intended classification task.

#### Developing mouse

We used a large-scale single-cell atlas of mouse development spanning embryonic and postnatal stages [14]. Gene names were mapped to Ensembl identifiers for compatibility with probe design resources. Stratified sampling was used to construct a balanced dataset of 500,000 cells based on the author-provided cell-type annotations (*K* = 190 categories).

#### Mouse lymph node

We constructed a composite lymph node reference by merging multiple published mouse lymph node scRNA-seq datasets (listed at [15–18]). Cell-type labels were harmonized across datasets, and doublets were removed using the original study annotations. The final dataset was stratified to 100,000 cells across 43 harmonized cell types.

### CIPHER Architecture

CIPHER is a neural-network architecture that couples a biologically constrained linear encoder to a nonlinear decoder. The encoder represents the experimental design, that is, how probes are allocated across genes and readout channels, whereas the decoder evaluates whether the resulting projected measurements preserve cell-type information.

#### Encoder (experimental design layer)

The encoder learns an encoding matrix *E* ∈ ℝ ^*G×B*^, where *G* is the number of genes and *B* is the number of bits. Each entry *E*_*ij*_ represents the number of probes assigned from gene *i* to bit *j*. To ensure that probe allocations are nonnegative and bounded by the number of available probes for each gene, the trainable encoder parameters *W*_*ij*_ are transformed as

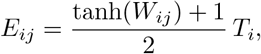

where *T*_*i*_ is the maximum number of probes available for gene *i*. Thus, each encoder entry is constrained to the interval [0, *T*_*i*_]. The values of *T*_*i*_ were fixed in advance from probe-design feasibility and depended only on gene-specific properties such as transcript length and probe availability in PaintSHOP [19]. During optimization, *E* is treated as continuous; for probe synthesis, the final optimized matrix is discretized by rounding and then clipped to satisfy the per-gene constraints exactly.

Given a normalized expression matrix *X* ∈ ℝ^*N ×G*^ for *N* cells, the encoder defines the latent projection

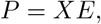

where *P* ∈ ℝ^*N ×B*^ is the predicted bit-level signal matrix. In this formulation, each bit is a continuous-valued aggregate transcriptional measurement obtained by summing contributions from multiple genes rather than a binary on or off code.

#### Decoder (cell-type prediction layer)

The decoder maps the projected signals *P* to cell-type predictions. It is implemented as a multilayer perceptron with *h* hidden layers, each of width 3*B*. Each hidden layer consists of a linear transformation followed by batch normalization and a GELU nonlinearity. The final layer is linear and outputs class logits, which are converted to class probabilities by the cross-entropy loss during training. Unless otherwise noted, all experiments used a single hidden layer (*h* = 1).

Before decoding, projected signals can optionally be normalized within each cell by their total projected intensity and, if desired, standardized across bits. In the experiments reported here, the decoder operated on the projected bit intensities directly unless otherwise stated. The decoder therefore serves only as a differentiable objective for evaluating whether a proposed probe allocation preserves cell-type information; it is not part of the experimental measurement itself.

#### Initialization and design interpretation

Encoder weights were initialized so that, after the bounded transformation above, the initial probe fractions for each gene–bit pair lay within a user-specified interval. This avoids starting from either nearly empty or fully saturated probe allocations. The learned encoding matrix can be interpreted directly as an experimental design: each column defines one bit, and the entries in that column specify how probe counts are distributed across genes to produce that aggregate readout channel.

### Loss Functions and Optimization

CIPHER is trained end-to-end by minimizing a weighted objective with four conceptual components:

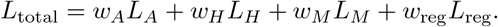

For clarity, we present the objective in terms of accuracy, hybridization, measurability, and optional regularization components, although in implementation each constituent term has its own user-defined coefficient. In the experiments reported here, the active optimization terms were *L*_*A*_, *L*_*H*_, and *L*_*M*_.

#### Accuracy loss (*L*_*A*_)

Let *P* = *XE* denote the latent projection of the normalized expression matrix *X* through the encoding matrix *E*, and let *f*_*θ*_(*P*) denote the decoder output. Cell-type discrimination was enforced using categorical cross-entropy with label smoothing:

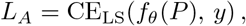

where *y* is the ground-truth cell-type label and the label-smoothing coefficient was fixed at *ϵ* = 0.1.

#### Hybridization constraints (*L*_*H*_)

The hybridization term combines a global probe-budget constraint and a per-gene probe-allocation constraint:

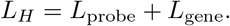

For the global budget, let

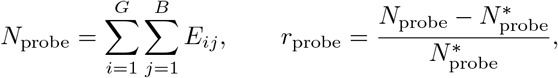

where 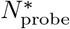 is the target total number of probes. The implementation applies a soft penalty to the relative excess probe count using an ELU nonlinearity:

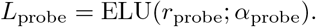

For the per-gene constraint, let *T*_*i*_ denote the maximum number of probes available for gene *i*, and define the excess allocation

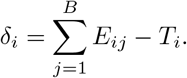

Only genes exceeding the available probe budget contribute to the loss:

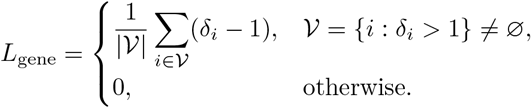

#### Measurability losses (*L*_*M*_)

The measurability term combines brightness, dynamic range, and cell-type separation:

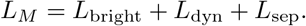

These quantities are computed from clean latent projections after per-cell sum normalization. Let *P*_*cb*_ denote the projected signal of cell *c* in bit *b*, and define

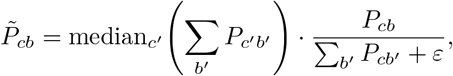

where *ε* = 10^−8^ prevents numerical instability. For each bit *b*, we then define three summary statistics from quantile windows of 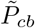 across cells:

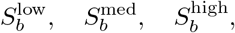

corresponding to the mean signal over the 5th–15th, 45th–55th, and 85th–95th percentile intervals, respectively. Brightness penalizes bits whose median signal falls below a target brightness *S*^*^:

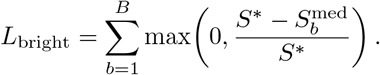

Dynamic range is defined for each bit as

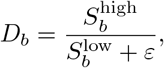

and bits whose dynamic fold falls below the target *D*^*^ are penalized:

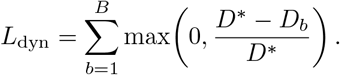

Cell-type separation is computed from cell-type centroids in normalized latent space. Let *v*_*k*_ ∈ ℝ^*B*^ denote the mean normalized bit profile of cell type *k*. For each pair of cell types (*k, ℓ*), separation is defined as the largest relative bit-wise difference:

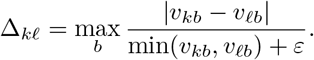

Pairs whose separation falls below the target Δ^*^ are penalized according to

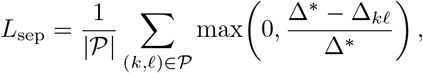

where 𝒫 is the set of cell-type pairs represented in the current batch.

#### Training schedule and optimization

Unless otherwise stated, models were trained with *B* = 18 bits and a single decoder hidden layer. Optimization used Adam with mini-batches of 500 cells for 100,000 iterations. Several target parameters, including brightness, dynamic fold, separation fold, and learning rate, were linearly interpolated between user-specified initial and final values during training. If a parameter *θ* had start and end values *θ*_*s*_ and *θ*_*e*_, its value at iteration *t* was

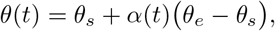

with

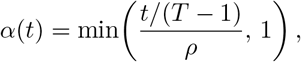

where *T* is the total number of training iterations and *ρ* is the saturation fraction controlling when the final target is reached. Unless otherwise specified, key targets were ramped over the first 75% of training. The objective is non-convex because it combines a constrained nonlinear encoder–decoder architecture with stochastic optimization and multiple soft constraint terms.

### Selection of Target Values

In practice, we parameterized CIPHER by anchoring the target goals to simple assay-level estimates rather than treating them as abstract optimization knobs. The central quantity is the expected signal generated by a bit. For a given bit *b*, the expected signal can be approximated as

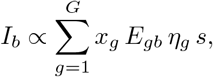

where *x*_*g*_ is the transcript abundance of gene *g, E*_*gb*_ is the allocated probe count from gene *g* to bit *b, η*_*g*_ is an effective probe-to-signal conversion factor that subsumes hybridization and detection efficiency, and *s* is the microscope sensitivity expressed in camera units per detected fluorophore. This expression was used only as a practical guide for selecting reasonable target values, not as an exact physical forward model.

In our imaging system, a single Cy5 fluorophore corresponded approximately to 0.5 camera arbitrary units, and in practice we considered signals in the high hundreds of camera units to be comfortably above the empirical detection floor. To estimate the effective conversion factor *η*_*g*_, we used literature values as an order-of-magnitude guide. In particular, bDNA-amplified MERFISH measurements have been reported to achieve hybridization efficiencies on the order of 60% under favorable conditions [20]. We therefore treated the overall probe-to-signal conversion as substantially below unity and used this estimate only to place the brightness target in a realistic regime. The resulting brightness target was intended to place the median bit signal safely above the noise floor without forcing the design toward arbitrarily bright but weakly discriminative solutions.

The total probe budget 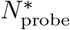 was chosen jointly with the brightness target because achievable bit brightness depends both on transcript abundance and on the number of accessible probes per transcript. Increasing the probe budget can increase bit intensity, but only within the limits imposed by transcript structure, probe-design feasibility, and practical library size. In this sense, 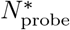 and *S*^*^ were treated as coupled experimental design parameters rather than independent optimization targets.

The dynamic-fold and separation-fold targets were selected to exceed the expected technical variability of the assay and to preserve margins between closely related cell types. Specifically, the dynamic-fold target

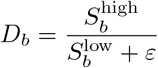

was set high enough that each bit retained useful contrast across the observed signal range, while the separation target

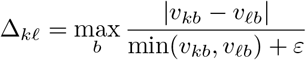

was chosen to preserve a minimum relative margin between cell-type centroids in projected space. In practice, noisier assays or finer-grained cell-type distinctions motivated larger values of *D*^*^ and Δ^*^, whereas cleaner assays or coarser classification tasks permitted weaker targets.

By contrast, the relative loss weights were used only to determine how strongly the optimizer prioritized these experimentally motivated target goals once they had been specified. Consistent with our sensitivity analysis, several-fold changes in the measurability loss weights altered the balance among robustness metrics more than they altered overall performance, so in most applications we recommend selecting biologically realistic target values first and leaving the default weights unchanged unless a specific unmet objective needs to be emphasized.

### Noise Injection and Robustness

Noise was introduced at two distinct stages: as training-time augmentation within CIPHER and as a post hoc simulation benchmark used to compare learned encodings under measurement perturbations.

#### Training-time noise

During optimization, noise was applied in latent space to stabilize training and to model nonzero experimental background. Let

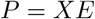

denote the clean latent projection. During training, the decoder received a perturbed version

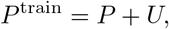

where

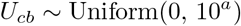

independently for each cell *c* and bit *b*, and *a* is a user-defined additive background parameter. In the default training configuration used here, this term was the only active noise process. Although the implementation also supports optional gene-level multiplicative perturbations, encoder-weight perturbations, projection-level multiplicative noise, and dropout, these additional noise terms were assigned zero weight in the experiments reported in this study.

#### Simulation-based robustness benchmark

To evaluate robustness of learned encodings under more realistic measurement variability, we performed a separate simulation analysis after training. In this benchmark, each projected bit value was perturbed by signal-dependent multiplicative noise rather than a fixed additive offset. For a simulated projected matrix *X* ∈ ℝ^*N ×B*^, noisy measurements were generated as

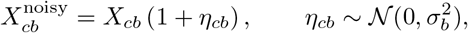

followed by clipping to nonnegative values:

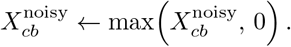

The bit-specific noise scale *σ*_*b*_ was calibrated from the nominal benchmark noise level *λ* by matching the variance of the previous additive perturbation at the median signal level of each bit. Let

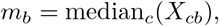

with

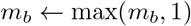

to avoid numerical instability for very weak bits. Since a uniform perturbation on [0, *λ*] has standard deviation 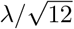, the corresponding multiplicative noise scale was chosen as

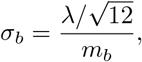

and then clipped to the interval [0, 0.5]:

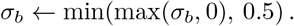

This calibration makes the perturbation magnitude easy to control in a way that remains interpretable across designs. For each nominal noise setting, the corresponding multiplicative noise level is chosen so that the induced variation at the median signal level of each bit matches a specified reference scale. As a result, weak bits experience larger relative perturbations than strong bits, which is consistent with signal-dependent measurement uncertainty, while the overall severity of the perturbation remains easy to tune through a single scalar setting. This provides a simple way to increase or decrease noise strength without having to choose separate multiplicative parameters for each bit by hand.

### Evaluation Metrics

Performance was evaluated on held-out test data using classification accuracy, separation, and Dynamic Fold, all computed from the clean latent projections on the test set. Accuracy was defined as the fraction of test cells whose predicted label matched the ground-truth label:

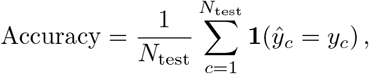

where *ŷ*_*c*_ is the decoder prediction for cell *c* and *y*_*c*_ is the corresponding reference label.

The separation and Dynamic Fold metrics were computed from the same normalized latent representation used in the measurability losses. Let *P*_*cb*_ denote the clean projected signal of cell *c* in bit *b*, and define the per-cell normalized bit signal

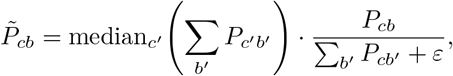

with *ε* = 10^−8^.

For each bit *b*, we defined three summary statistics from quantile windows of 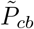 across all test cells:

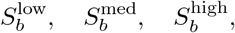

corresponding to the mean signal within the 5th–15th, 45th–55th, and 85th–95th percentile intervals, respectively. The Dynamic Fold of bit *b* was then defined as

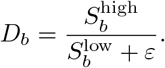

The reported *Dynamic Fold* metric was the 10th percentile of 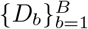 across bits:

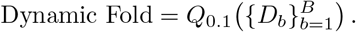

Using the 10th percentile rather than the mean emphasizes the lower tail of bit performance and penalizes designs in which a subset of bits is weak or poorly distributed.

To quantify cell-type separation, we first computed the centroid of each cell type in normalized latent space. Let 𝒞_*k*_ denote the set of test cells assigned to cell type *k*, and define the centroid

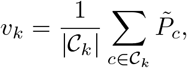

where 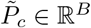 is the normalized bit vector for cell *c*. For each pair of cell types (*k, ℓ*), separation was defined as the largest relative bit-wise difference between their centroids:

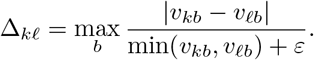

The reported *Separation* metric was the 10th percentile of Δ_*kℓ*_ across all cell-type pairs present in the test set:

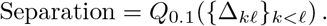

This summary again emphasizes lower-tail behavior and therefore captures whether a design leaves a substantial fraction of cell-type pairs only weakly separated, rather than reporting only the average or best-resolved distinctions.

In addition to these summary metrics, the evaluation code also recorded the corresponding no-noise, low-noise, medium-noise, and high-noise accuracies, as well as auxiliary quantities such as Dynamic Range and per-bit signal percentiles. Unless otherwise stated, the main performance values reported in the manuscript refer to the clean test-set metrics defined above.

### Sensitivity to loss weighting

To assess whether CIPHER depended strongly on the precise relative weighting of the measurability objectives, we performed a sensitivity analysis in which the coefficient of one measurability term was varied at a time while all other model settings were held fixed. Separate sweeps were performed for the brightness, dynamic-range, and separation terms. For each setting, models were retrained from the same reference dataset and evaluated on held-out test cells using the standard prediction accuracy, separation, and Dynamic Fold metrics defined above.

### Cross-chemistry transfer test

To evaluate sensitivity to reference shift, we exploited the separation between 10x v2 and v3 chemistries within the Allen whole-brain reference dataset. Using the same preprocessing and gene set shared across chemistries, CIPHER was trained separately on the v2 and v3 subsets and then evaluated on held-out test cells from both the matched and alternate chemistry, yielding the four train-test combinations v2 → v2, v2 → v3, v3 → v3, and v3 → v2. This analysis was used as a controlled stress test of chemistry-dependent reference variation.

### GO term enrichment

To assess the biological interpretability of the learned aggregate features, we performed Gene Ontology enrichment analysis separately for each bit of the discretized encoding matrix. For each bit, we defined the query set as all genes with non-zero weight in that column of the constrained encoding matrix and used the full set of genes present in the matrix as the background. Enrichment was computed with g:Profiler (`gprofiler`) for `GO:BP, GO:MF`, and `GO:CC` terms using the *Mus musculus* annotation, and terms were retained using false discovery rate correction with a significance threshold of 0.05.

### Probe Sequence Design and Assembly

Encoding matrices were converted into oligonucleotide libraries using PaintSHOP-designed 30-mer gene-specific probes [19]. For each gene–bit pair, integer probe counts were allocated according to the discretized encoding matrix. Readout sequences were drawn from a prevalidated orthogonal set and assembled into final probe constructs using rolling circle amplification (RCA). Full experimental synthesis details will be provided elsewhere.

### Experimental Imaging and Decoding

ATLAS experiments were performed on fresh-frozen adult mouse brain coronal sections using CIPHER-designed probe pools and automated cyclic fluorescence imaging, as described previously for the ATLAS platform. Briefly, sections were hybridized with large oligonucleotide probe libraries encoding aggregate transcriptional signatures, imaged over repeated rounds of readout hybridization and stripping, and segmented to obtain single-cell bit-intensity profiles across whole-brain sections. The resulting bit-level measurements were decoded with the SCALE framework, which aligns measured signals to scRNA-seq references through hierarchical classification and spatial priors defined in the Allen Common Coordinate Framework. Predicted cell-type labels were then used to impute full transcriptomes by nearest-neighbor averaging within the aligned latent space. In the present study, these experimental procedures were unchanged from our previous ATLAS implementation; only the design of the aggregate probe panels was modified by the CIPHER optimization framework. Full details of tissue processing, probe hybridization, cyclic imaging, image registration, segmentation, decoding, and transcriptome imputation are provided in our previous ATLAS study. [12]

## Results

CIPHER (Cell Identity Projection using Hybridization Encoding Rules) is a neural-network framework for the joint design and decoding of transcriptional signature measurements (Figure 1). The model links single-cell RNA sequencing (scRNAseq) expression profiles (*X*) to cell-type identities (*C*) through an intermediate latent representation (*P*) that defines a low-dimensional projection of transcriptional space. Each dimension of *P* corresponds to an aggregate transcriptional signature, implemented as a weighted combination of genes that can, in principle, be realized *in situ* using pools of DNA oligonucleotide probes, hereafter referred to as encoding probes. Unlike prior aggregate encoding approaches that focus primarily on representing reference information accurately, CIPHER explicitly incorporates the experimental realizability of these signatures. Its objective is to design transcriptional signatures that remain maximally informative under realistic molecular and optical constraints imposed by *in situ* hybridization within intact tissue.

CIPHER adopts a neural-network architecture that accommodates multiple design objectives through distinct loss functions (*L*_*H*_, *L*_*M*_, and *L*_*A*_; Figure 1). The encoder constitutes the first linear layer of the network and represents the molecular design of the experiment, specifying how many probes hybridize to each gene and how their combined signals contribute to each readout channel. Because probe hybridization is additive and implemented through physical oligonucleotides, encoder weights are constrained to be nonnegative and correspond to the number of encoding probes assigned per gene–channel pair. This layer performs a linear projection from gene space to latent space, directly mirroring the physical encoding of gene combinations *in situ*.

The decoder introduces nonlinearity through hidden layers that map transcriptional signatures to predicted cell types. Unlike the encoder, the decoder has no molecular counterpart and exists purely *in silico*. Together, these components capture the linear physics of measurement and the nonlinear decision boundaries of cell-type classification within a unified framework. Training is performed end-to-end using scRNA-seq data, allowing the encoder and decoder to co-optimize such that the learned encoding matrix reflects both biological structure and experimental constraints.

CIPHER incorporates three complementary loss terms, each targeting a distinct design objective. The hybridization loss *L*_*H*_ penalizes excessive probe usage per gene to promote sparse, experimentally feasible encodings. A weight *w*_*ij*_ mapping gene *i* to readout channel *j* corresponds to the number of bivalent encoding probes hybridizing to gene *i* and contributing to channel *j*. Because the number of accessible binding sites on an mRNA is finite and depends on transcript length and sequence composition, the sum of *w*_*ij*_ across channels is physically constrained.

The accuracy loss *L*_*A*_ enforces biological fidelity by measuring categorical cross-entropy between predicted and known cell-type labels derived from the scRNA-seq reference. This term captures computational decoding performance independent of experimental considerations. The measurability loss *L*_*M*_ promotes robustness under realistic experimental conditions by encouraging broad separation of cell-type signatures in signal space and sufficient dynamic range of encoded features. This term avoids degenerate solutions in which cell types are separable in principle but differ only weakly in absolute signal, rendering them difficult to distinguish experimentally. Together, these losses balance biological informativeness, computational accuracy, and experimental feasibility within a unified optimization framework.

We next examined how encoding dimensionality and loss-function composition influence CIPHER performance and latent-space structure (Figure 2a). All analyses used the Allen Mouse Brain Cell Type Atlas [13], which was also employed for ATLAS signature design. We trained CIPHER across increasing numbers of latent dimensions while systematically varying which loss terms were included.

Decoding accuracy increased rapidly with dimensionality before plateauing, indicating that most biologically relevant variation can be captured within a compact encoding space (Figure 2b). Models trained using only *L*_*A*_ achieved high accuracy with as few as 10–15 latent features. Adding the sparsity constraint *L*_*H*_ caused a modest reduction in accuracy, while inclusion of *L*_*M*_ produced an additional but small decrease. Using either component of *L*_*M*_ alone—dynamic range 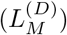 or separability 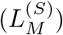—had a similar impact on accuracy as using both, indicating largely overlapping effects. These results show that incorporating experimental constraints does not substantially compromise decoding accuracy and that accuracy alone saturates early.

We quantified how each optimization objective affects cell-type separation in latent space (Figure 2c). Separation reflects robustness to measurement perturbations rather than idealized decoding accuracy. Models trained using only *L*_*A*_, with or without *L*_*H*_, achieved high accuracy but low separation, revealing a mismatch between *in silico* optimization and *in situ* measurement constraints. Inclusion of *L*_*M*_ directly addressed this gap: 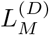 expanded dynamic range, 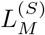 increased inter–cell-type distances, and combining both yielded the largest gains, particularly at higher dimensionality. Dynamic fold exhibited the same behavior, remaining narrow under *L*_*A*_ and *L*_*H*_ alone and expanding substantially only when *L*_*M*_ was included. These gains occurred with only minor reductions in accuracy, demonstrating that accuracy alone is insufficient as a design criterion.

To assess the nonlinear capacity required for decoding, we varied the depth of the decoder *f*_*D*_ while holding the encoder and loss configuration fixed (Figure 3a). A purely linear decoder (*h* = 0) underperformed at low dimensionality, particularly with 3–6 features, indicating that a strictly linear mapping from *P* to *C* is insufficient. Introducing a single hidden layer largely eliminated this gap, while deeper decoders provided no consistent benefit (Figure 3). Accordingly, all subsequent analyses use a decoder with one hidden layer (*h* = 1).

**Fig 3.**
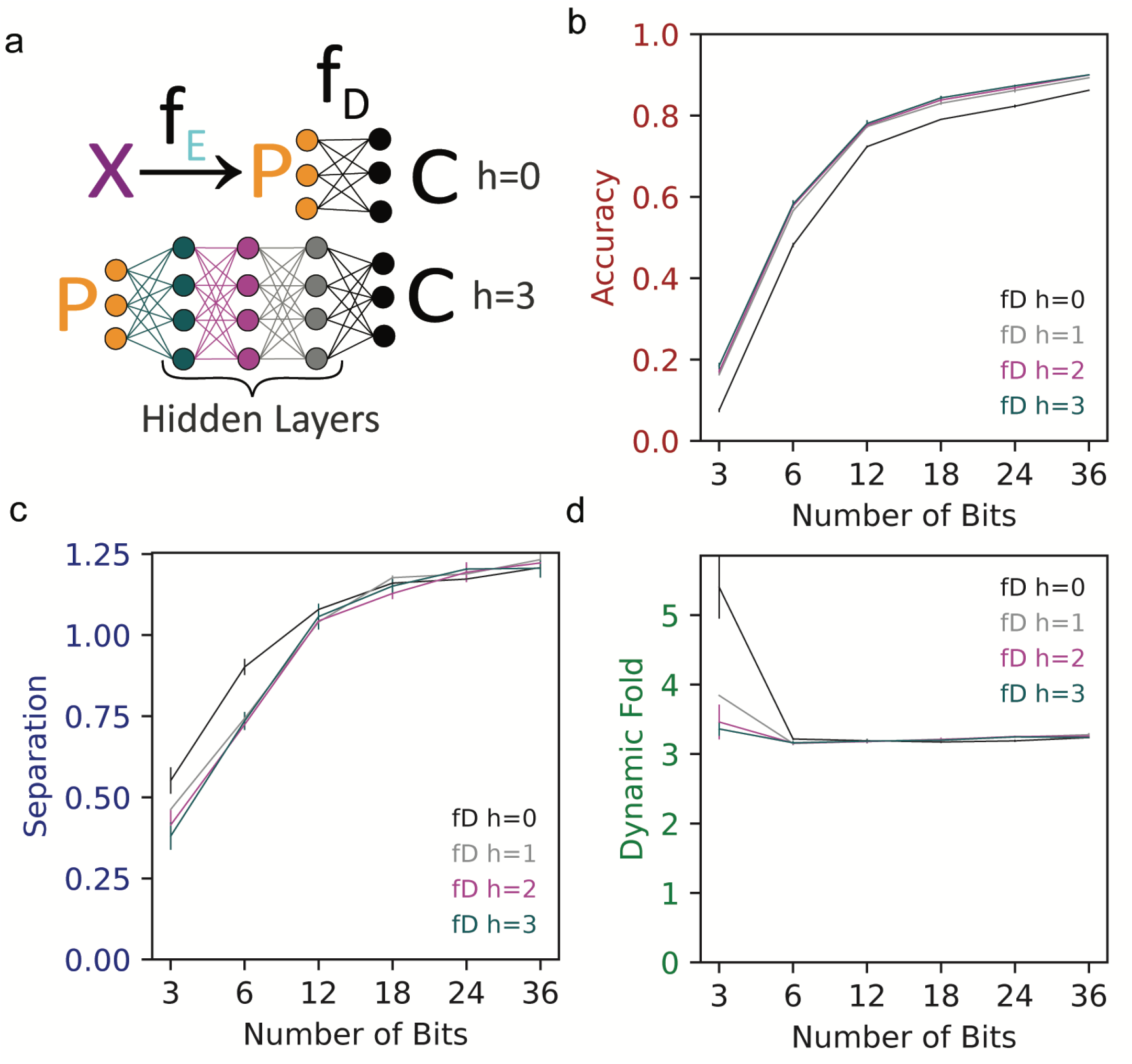
Linear vs. Nonlinear Decoder Complexity. (A) Schematic comparing decoder architectures with varying depths (number of hidden layers, *h*). A linear decoder (*h* = 0) represents simple weighted summation, whereas nonlinear decoders capture combinatorial patterns in the latent space. (B–D) Performance metrics plotted against the number of bits (imaging rounds) for different decoder complexities. (B) Classification accuracy increases with bit number but plateaus differently depending on decoder complexity. While performance improves modestly from a linear decoder (*h* = 0) to a shallow nonlinear decoder (*h* = 1), further increases in depth (*h >* 1) provide no meaningful additional gain. (C) Latent-space separability metrics. (D) Relationship between available bits and the required dynamic fold change, illustrating the trade-off between decoder complexity and signal constraints. See Methods for the definitions of Accuracy, Separation, and Dynamic Fold.

We next examined how signal-brightness constraints influence encoding robustness (Figure 4a). Relaxing brightness constraints caused a sharp drop in separation, whereas accuracy and dynamic fold declined only modestly. This indicates that brighter encodings often collapse inter–cell-type distances by recruiting weakly discriminative genes, reducing robustness despite high decoding accuracy. In contrast, varying the total probe budget produced gradual and parallel effects across all metrics (Figure 4b), with performance saturating at approximately 50,000 probes, a regime compatible with current synthesis capabilities.

**Fig 4.**
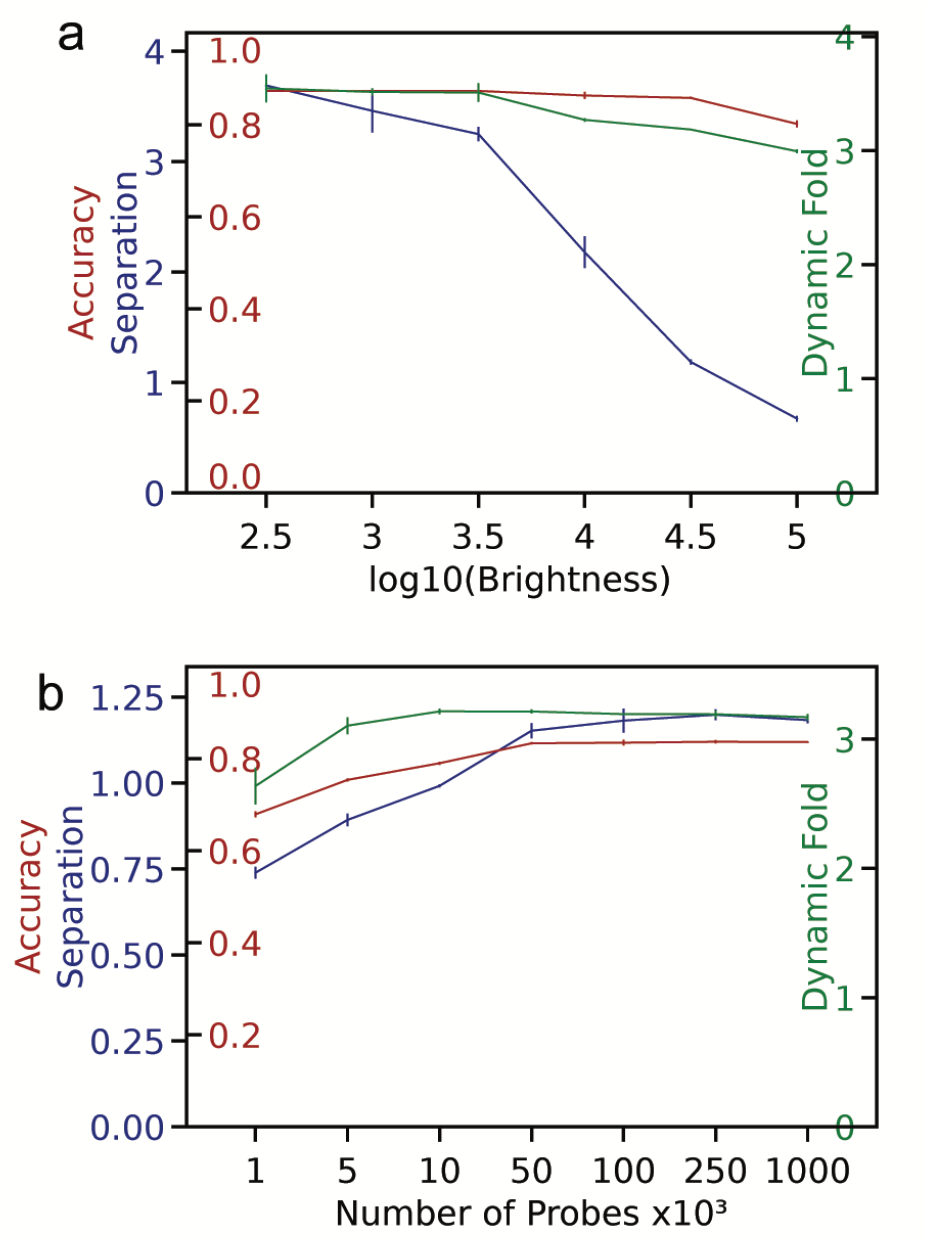
Experimental Constraints and Probe Design. (A) Encoding performance as a function of target signal brightness. The x-axis represents the target median brightness (log_10_ scale). The left y-axis displays latent-space separability (blue) and classification accuracy (red), while the right y-axis displays dynamic fold change (green). Results indicate a strong advantage to lower brightness requirements, which are ultimately determined by microscope sensitivity and hybridization efficiency. (B) Encoding performance as a function of total probe budget. The x-axis represents the maximum number of encoding probes (*×* 10^3^). Axes for separability (blue, left), accuracy (red, left), and dynamic fold (green, right) correspond to those in panel (A). Performance metrics plateau at approximately 5 *×* 10^4^ probes, indicating diminishing returns for larger probe libraries. See Methods for the definitions of Accuracy, Separation, and Dynamic Fold.

We next examined whether CIPHER performance was more sensitive to the precise weighting of its optimization terms or to the reference data used for design (Figure 5). Several-fold changes in the individual measurability weights produced only gradual changes across evaluation metrics. Increasing the separation term shifted the tradeoff toward larger latent-space margins and higher Dynamic Fold, with only a modest reduction in prediction accuracy, whereas varying the brightness and dynamic-range terms had comparatively small effects. These results indicate that CIPHER does not rely on narrow tuning of the measurability loss coefficients to achieve reasonable performance.

**Fig 5.**
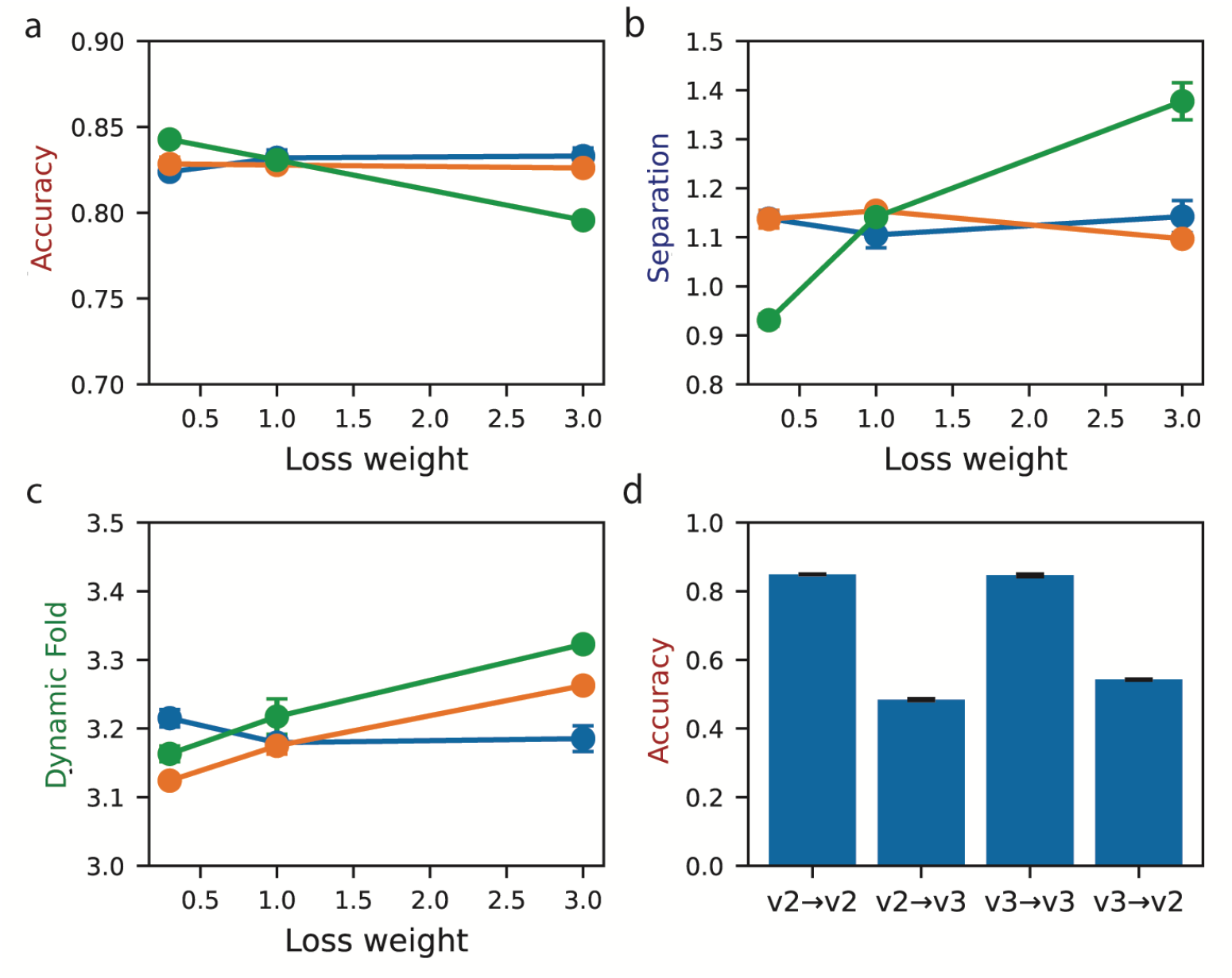
Sensitivity of CIPHER to loss weighting and reference shifts. (A–C) Performance metrics plotted against the weight of individual measurability terms. Separate sweeps were performed for the brightness, dynamic-range, and separation losses while all other model settings were held fixed. Panels show (A) prediction accuracy, (B) separation, and (C) Dynamic Fold on held-out test data. (D) Cross-chemistry transfer test using the 10x v2 and v3 partitions of the Allen whole-brain reference. Models were trained on one chemistry and evaluated on held-out cells from either the same or the alternate chemistry, yielding the four train-test combinations shown.

We then asked whether performance was more strongly affected by shifts in the reference dataset itself. To test this, we exploited the separation between 10x v2 and v3 chemistries within the Allen whole-brain reference and evaluated all four train-test combinations (Figure 5d). In contrast to the modest effects of loss reweighting, cross-chemistry transfer produced a substantially larger reduction in accuracy than within-chemistry evaluation. Thus, CIPHER is relatively stable to moderate changes in loss weighting but more sensitive to systematic differences in the reference data used for training and deployment.

We compared CIPHER to DPNMF, a supervised discriminant non-negative matrix factorization approach previously used in ATLAS, using matched dimensionality and the same scRNA-seq reference [21, 22]. Similar to CIPHER, DPNMF uses a supervised approach to identify latent dimensions and shares other important characteristics with CIPHER, such as sparsity and non-negativity. Furthermore, DPNMF was used in prior experimental work that uses transcriptional signatures, making it the appropriate comparison to CIPHER. DPNMF concentrates separability into coarse divisions, primarily neurons versus non-neurons, whereas CIPHER distributes separability across subclasses (Figure 6a). Correspondingly, DPNMF exhibited a long tail of poorly decoded subclasses (Figure 6b) and highly uneven brightness distributions dominated by a few saturated features (Figure 6c). CIPHER avoided these pathologies and maintained robustness under simulated noise perturbations (Figure 6d). To assess biological interpretability of the learned encodings, we examined the genes assigned to each CIPHER bit and summarized their associated functions by gene ontology enrichment analysis. Across bits, enriched terms were consistent with expected neuronal, glial, and regional transcriptional programs rather than arbitrary gene mixtures, supporting the biological plausibility of the learned aggregate signatures. The full gene and probe composition of each bit, together with enrichment results, is provided in Supplementary Table 1.

**Fig 6.**
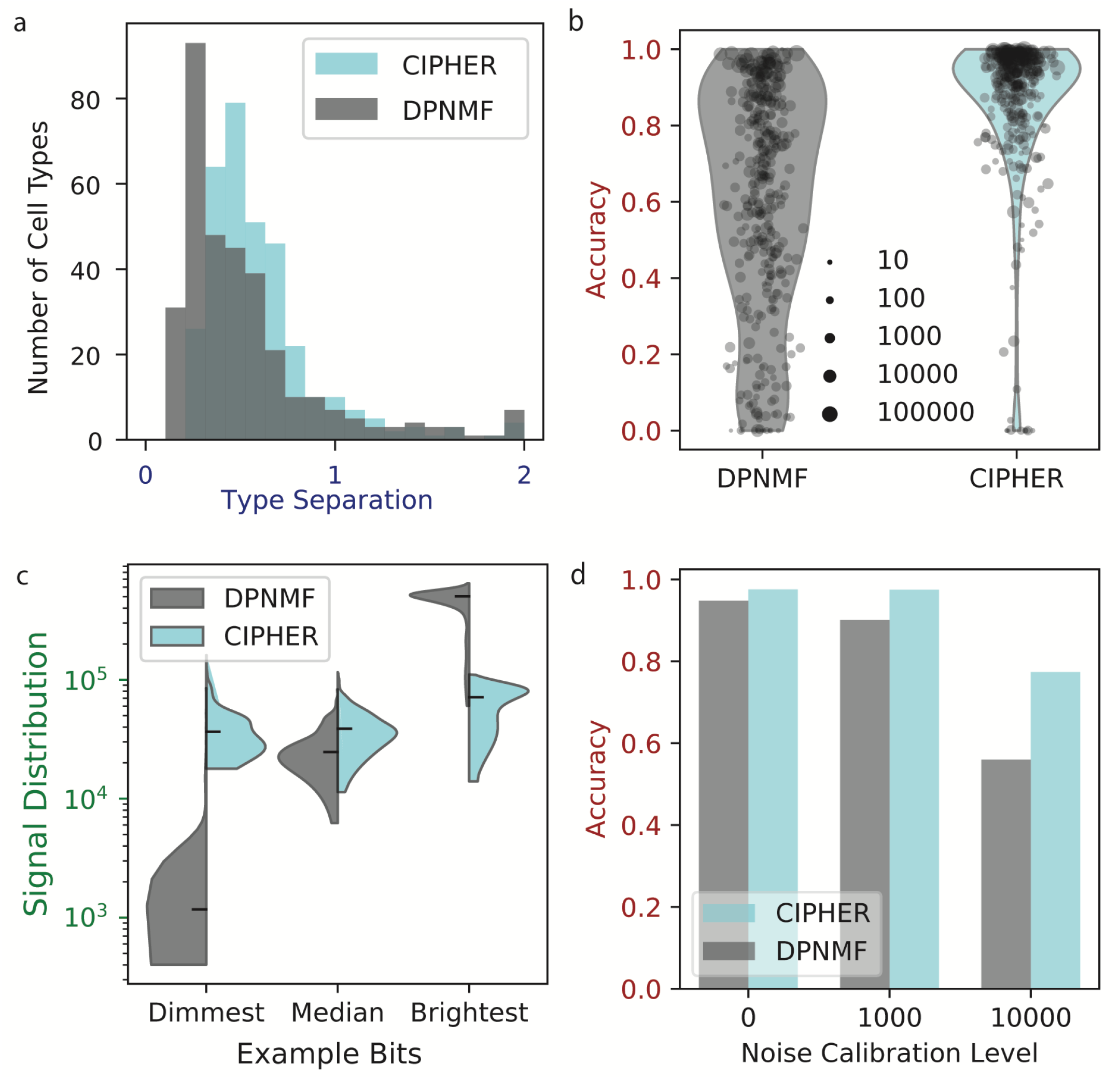
Comparison of CIPHER and DPNMF Performance. (A) Histogram of pairwise distances between cell-type centroids in the learned latent space. DPNMF encodings exhibit a larger fraction of cell types with near-zero separation compared to CIPHER. (B) Classification accuracy broken down by individual cell types in simulation, demonstrating that a substantially larger proportion of cell types achieve high accuracy with CIPHER than with DPNMF. (C) Distribution of individual bit brightness values. CIPHER encodings maintain relatively uniform signal brightness across bits, whereas DPNMF encodings display high variance with pronounced disparities between features. (D) Simulation using imputed MERFISH data showing expected classification accuracy under increasing levels of multiplicative noise. Noise levels were calibrated so that their overall effect matches 0,1000,100000 counts. CIPHER maintains higher accuracy and substantially greater robustness to noise than DPNMF.

We validated CIPHER *in situ* in mouse cortex, decoding cell types with SCALE and observing expected laminar and hippocampal organization (Figure 7). Greedy spatial matching to MERFISH data showed high concordance, providing an upper bound on achievable agreement given imperfect registration. Finally, we applied CIPHER to the developing mouse brain and lymph node (Figure 8), observing tissue-specific saturation of dimensionality at ∼ 24 and ∼ 9 features, respectively. These results demonstrate that CIPHER identifies compact, context-dependent encodings and generalizes across diverse biological systems.

**Fig 7.**
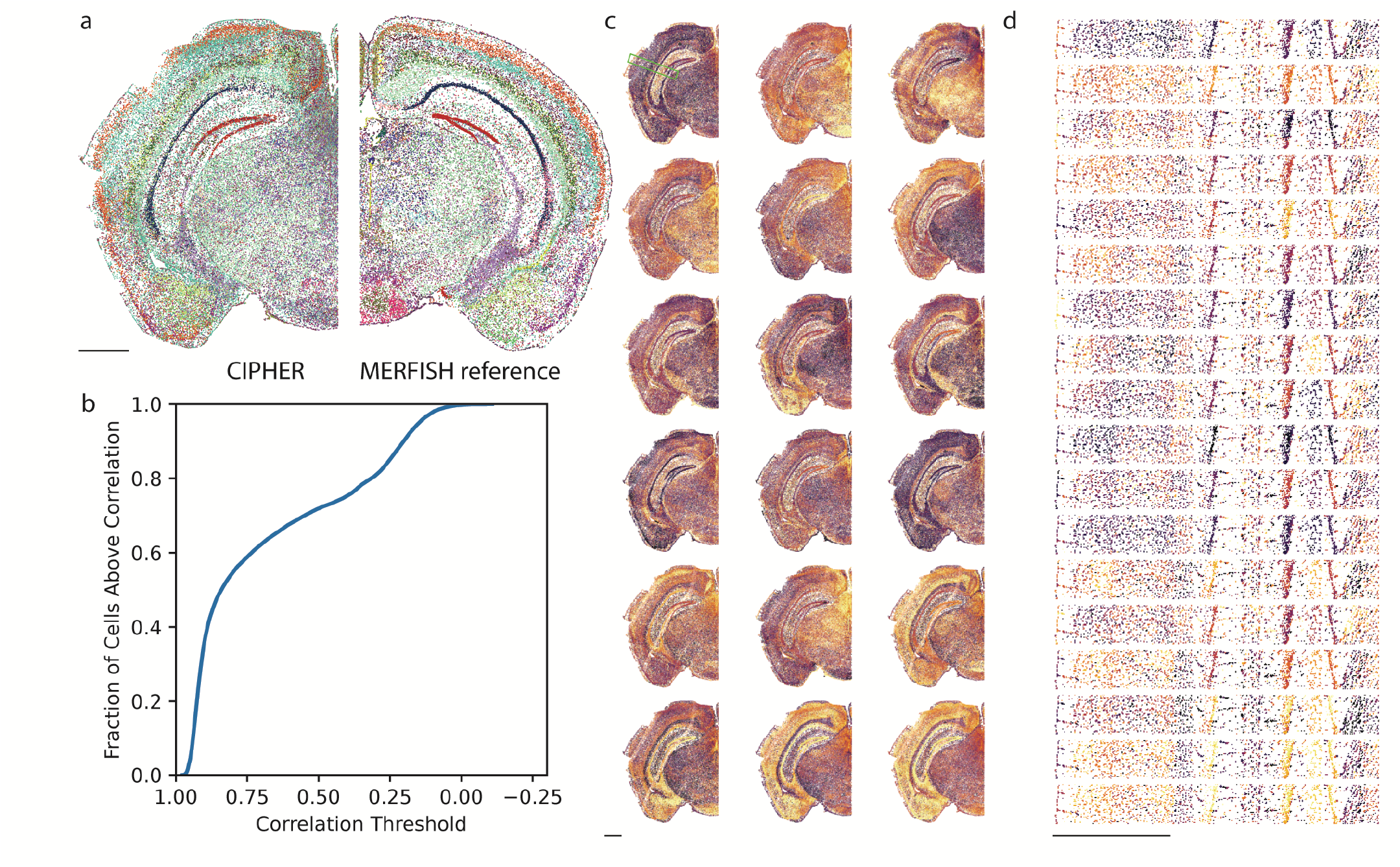
Experimental Validation in Mouse Cortex. (A) Left: Decoded cell-type map for a representative half-coronal mouse cortex section, where each point corresponds to a single cell colored by assigned identity. Right: Corresponding reference map derived from MERFISH data for the same anatomical region. Colors match BICCN cell type color mapping [13] (B) Concordance between CIPHER measurements and reference data. Cells were greedily paired across datasets based on spatial proximity, and agreement was quantified as the correlation between their gene expression profiles. Because the datasets are not registered at single-cell precision, this procedure provides an upper bound on achievable agreement within local neighborhoods. (C) Visualization of the measured latent-space signals across the tissue section. Heatmap intensities reflect bit values, with dimmer cells shown in purple and brighter cells in yellow, illustrating the combinatorial encoding. The green rectangle in the first image shows the location of the zoomed view shown in panel D. (D) High-resolution view highlighting cortical layers and hippocampal subfields (a zoomed view of panel C). Insets reveal individual cells and distinct combinatorial signal patterns. Scale bars in panels A-C are 1 mm.

**Fig 8.**
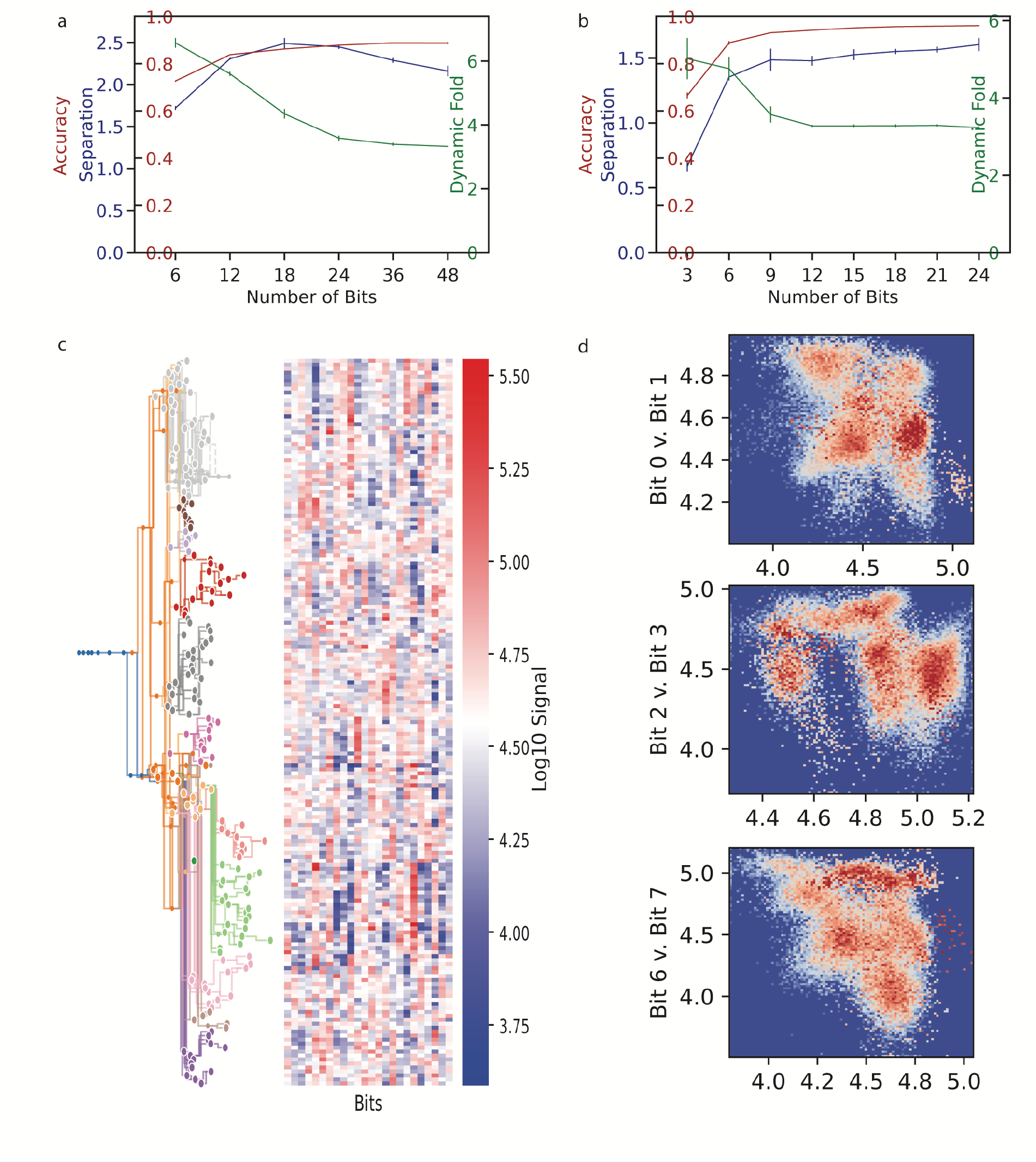
Generalization to Alternative Tissues. Demonstration of the CIPHER framework’s applicability across diverse biological contexts. (A) Performance metrics for the developing mouse brain plotted against the number of bits. Axes display separability (blue, left), accuracy (red, left), and dynamic fold (green, right). Performance saturates at approximately 24 bits. (B) Performance metrics for the lymph node plotted against the number of bits, showing saturation at approximately 9 bits. (C) Heatmap of learned latent vectors (*P*) organized by developmental lineage, illustrating structured assignment of progenitor and neuronal populations to specific bits. (D) Density visualization of the projection space for lymph node data, demonstrating clear separation between major immune cell populations across latent dimensions. The panel shows kernel density estimates of 2D slices of the overall probability density function.

## Discussion

Aggregate spatial transcriptomics methods redefine the goal of *in situ* measurement: rather than recovering complete transcriptomes gene by gene, they aim to acquire just enough structured information to reliably align spatial observations with rich scRNAseq references. This reframing has enabled dramatic gains in scale and throughput, but it also introduces a new and largely unresolved design problem. Once individual genes are no longer the fundamental measurement unit, the central question becomes how to construct aggregate features that are not only informative in transcriptional space but also robust to the physical constraints and noise inherent to *in situ* hybridization and imaging. Our results show that existing design strategies, which optimize decoding accuracy or reconstruction fidelity in isolation, leave substantial performance on the table. CIPHER addresses this gap by treating aggregate encoding as a joint optimization problem over biological informativeness, signal robustness, and experimental feasibility.

A central insight emerging from our analyses is that classification accuracy alone is an inadequate objective for aggregate encoding design. Across tissues and model configurations, decoding accuracy saturates rapidly with relatively few encoded features, often well before the encoding becomes robust to measurement variability. This behavior reflects a fundamental mismatch between in silico optimization and *in situ* data collection. When encodings are evaluated against the same scRNA-seq reference used for training, small margins between cell types are sufficient for high apparent accuracy. However, in physical experiments, hybridization efficiency varies across probes and genes, background fluorescence is spatially structured, and signal dynamic range is finite [23–25]. Under these conditions, narrow margins collapse quickly. By explicitly optimizing for latent-space separation and usable dynamic range, CIPHER exposes trade-offs that are invisible to accuracy-based optimization and demonstrates that modest sacrifices in nominal accuracy yield disproportionate gains in robustness. These results also suggest a practical interpretation for applying CIPHER across tissues and assay regimes. If designs achieve good nominal accuracy but poor robustness, increase separation and dynamic-range targets rather than decoder complexity. If bit intensities are systematically weak, increase brightness and probe-budget targets jointly. If performance saturates early with excess measurement capacity, reduce the number of bits or relax brightness targets.

These findings place CIPHER in a distinct position relative to existing aggregate spatial transcriptomics frameworks. CISI introduced the idea of composite measurements using compressed sensing, focusing on efficient reconstruction of individual gene expression profiles from random mixtures [10]. While powerful, this approach is agnostic to how discriminative power and signal strength are distributed across features, and does not explicitly constrain dynamic range or hybridization feasibility. FISHnCHIPs takes a different approach by grouping co-expressed genes into modules to amplify signal and improve sensitivity [11]. However, module selection remains largely heuristic and gene-centric, and the optimization is not performed end to end with the decoding task. ATLAS and related supervised factorization approaches explicitly align aggregate features with reference-defined cell types, enabling scalable atlas construction [12]. Yet, as our comparison to DPNMF, the encoding scheme used in ATLAS, illustrates, supervised alignment alone does not guarantee robust encodings: without constraints on brightness balance, separability, and dynamic range, discriminative power can concentrate into a small number of saturated features, leaving fine-grained distinctions fragile.

By contrast, CIPHER integrates these considerations into a single framework. The encoder represents the physical design of the experiment, constrained by probe budgets and brightness limits, while the decoder captures the nonlinear decision boundaries required for cell-type assignment. Importantly, we find that this decoder need not be deep or complex. A single nonlinear hidden layer is sufficient to exploit the structure organized by the encoder, and deeper networks provide no consistent benefit. This observation underscores that the primary challenge in aggregate spatial transcriptomics lies not in classifier expressivity but in how information is projected into a measurable space. Once that projection is well designed, decoding becomes straightforward.

Our analysis of brightness and probe-budget constraints clarifies a critical asymmetry in aggregate encoding design. Aggressively increasing signal brightness produces encodings that remain accurate under idealized conditions but exhibit a sharp collapse in latent-space separability, rendering them highly sensitive to experimental noise. In contrast, constraining the total number of probes imposes a more uniform and predictable cost across accuracy, separation, and dynamic range, with performance plateauing at probe counts that are already experimentally accessible. These results identify brightness, rather than probe availability, as the dominant bottleneck in robust aggregate measurements. Importantly, they also suggest a practical path forward: combining CIPHER-designed encodings with post-hybridization signal amplification strategies, such as hybridization chain reaction (HCR), branched DNA amplification, or rolling-circle–based methods, should substantially improve separability and robustness without sacrificing encoding structure [26–29]. Because these amplification approaches increase signal uniformly across encoded features rather than redistributing gene weights, they are well matched to CIPHER’s emphasis on balanced signal allocation and margin preservation. This combination is expected to outperform naive brightness-driven panel design strategies by removing brightness as a design constraint altogether. With amplified readout, signal detectability no longer competes with separability, allowing aggregate encodings to achieve markedly higher and more uniformly distributed cell-type separation that remains robust across tissues, scales, and experimental conditions.

The comparison to DPNMF, the encoding used in ATLAS [12], illustrates the consequences of ignoring these constraints. Although DPNMF captures broad transcriptional structure, it allocates most separability to coarse divisions, such as neurons versus non-neurons, while leaving closely related subclasses weakly separated. This behavior is not incidental but follows directly from optimizing global reconstruction without regard to how separability and signal are distributed. As a result, DPNMF encodings exhibit highly uneven brightness profiles and degrade rapidly under simulated measurement noise. CIPHER, by explicitly enforcing balanced signal utilization and separation margins, produces encodings that maintain subclass-level resolution and degrade gracefully as noise increases. Notably, in relatively forgiving experimental contexts such as adult mouse cortex, both designs achieve acceptable decoding accuracy *in situ*. This observation is important: CIPHER does not claim universal superiority in all regimes. Rather, its advantages become most pronounced as measurement conditions become more challenging or as finer-grained distinctions are required.

Our *in situ* validation confirms that CIPHER-designed encodings can be implemented at scale and decoded reliably using existing pipelines [12]. The agreement with MERFISH-derived references, assessed through spatially local matching, demonstrates that CIPHER measurements capture local transcriptional structure with high fidelity. At the same time, this comparison highlights a broader issue shared by all spatial transcriptomics technologies: precise one-to-one cell correspondence across modalities is rarely achievable, and validation necessarily relies on neighborhood-level or distributional agreement [30, 31]. CIPHER does not resolve this limitation, but it ensures that the information captured by aggregate measurements is structured in a way that maximizes what can be reliably recovered given these constraints.

Extending CIPHER to mouse development dataset and the lymph node reveals an additional and conceptually important result: the dimensionality required for robust encoding is strongly tissue dependent. Developmental systems with continuous trajectories and overlapping transcriptional programs require substantially more encoded features to stabilize separation and dynamic range than lineage-structured immune tissues. This observation suggests that the number of required aggregate features reflects an intrinsic property of the biological system and the question being asked, rather than a fixed technological parameter. In this sense, CIPHER provides a quantitative way to estimate the effective dimensionality of spatial variation that must be measured to achieve robust integration with single-cell references.

Several limitations of the current framework warrant discussion. First, CIPHER depends on the availability and quality of scRNA-seq references and their associated taxonomies. If rare or novel cell states are absent from the reference, or if the taxonomy itself is incorrect, the encoding will faithfully optimize for the wrong distinctions. A controlled cross-chemistry stress test using the 10x v2 and v3 partitions of the Allen reference further showed that transfer across chemistries was substantially weaker than within-chemistry performance, indicating that CIPHER is sensitive not only to the biological representativeness of the reference atlas but also to technical shifts between training and deployment data. This limitation is consistent with the broader problem of batch effects in scRNA-seq, where differences in chemistry, protocol, and dataset composition can distort alignment, label transfer, and downstream classification even when the underlying cell populations are similar [32]. Second, the measurement model incorporated through the *L*_*M*_ loss captures coarse constraints on dynamic range and separability but does not account for sequence-specific probe effects, channel cross-talk, or spatially structured background. Incorporating richer, modality-specific noise models is an important direction for future work. Third, the framework is optimized for discrete cell-type classification and does not explicitly preserve continuous gradients or trajectories, which may be central in developmental or disease contexts [14]. This creates an inherent tradeoff. Encodings optimized for robust assignment to known classes may be less suitable for unsupervised discovery of rare, novel, or continuously varying states, especially when the reference taxonomy is incomplete or noisy. Nevertheless, aggregate measurements remain highly valuable when the main goal is whole-tissue profiling at scale with reliable alignment to a reference. Extending the loss functions to protect manifold structure rather than class margins alone would broaden the applicability of the approach. Finally, although we demonstrate compatibility with SCALE decoding, the encoder is trained with a specific decoder architecture, and formal guarantees of transfer across decoding algorithms remain an open question.

Despite these limitations, CIPHER establishes a principled foundation for measurability-aware design in aggregate spatial transcriptomics. By exposing the trade-offs between accuracy, robustness, and experimental feasibility, it shifts panel design from heuristic marker selection toward quantitative optimization grounded in physical constraints. More broadly, the framework suggests a path toward information-theoretic approaches to spatial measurement, in which the capacity of an assay is matched to the intrinsic structure of the biological system being studied. As spatial transcriptomics continues to scale across tissues, conditions, and organisms [1, 2], such principled design strategies will be essential for turning raw throughput into reliable biological insight.

## Acknowledgments

This project was supported by NIH grant R01-HG012925 to RW and JJL. The funders had no role in study design, data collection and analysis, decision to publish, or preparation of the manuscript.

## References

1. Marx V. Method of the Year: spatially resolved transcriptomics. Nat Methods. 2021;18(1):9–14.

2. Moses L, Pachter L. Museum of spatial transcriptomics. Nat Methods. 2022;.

3. Chen KH, Boettiger AN, Moffitt JR, Wang S, Zhuang X. Spatially resolved, highly multiplexed RNA profiling in single cells. Science. 2015;348(6233):aaa6090.

4. Eng CHL, Lawson M, Zhu Q, Dries R, Koulena N, Takei Y, et al. Transcriptome-scale super-resolved imaging in tissues by RNA seqFISH. Nature. 2019;568(7751):235–239.

5. Codeluppi S, Borm LE, Zeisel A, La Manno G, van Lunteren JA, Svensson CI, et al. Spatial organization of the somatosensory cortex revealed by osmFISH. Nat Methods. 2018;15(11):932–935.

6. He S, Bhatt R, Brown C, Brown EA, Buhr DL, Chantranuvatana K, et al. High-plex imaging of RNA and proteins at subcellular resolution in fixed tissue by spatial molecular imaging. Nat Biotechnol. 2022;40(12):1794–1806.

7. Plummer JT, Dezem FS, Cook DP, Park J, Zhang L, Liu Y, et al. Standardized metrics for assessment and reproducibility of imaging-based spatial transcriptomics datasets. Nat Biotechnol. 2025; p. 1–13.

8. Ståhl PL, Salmén F, Vickovic S, Lundmark A, Navarro JF, Magnusson J, et al. Visualization and analysis of gene expression in tissue sections by spatial transcriptomics. Science. 2016;353(6294):78–82.

9. Rodriques SG, Stickels RR, Goeva A, Martin CA, Murray E, Vanderburg CR, et al. Slide-seq: A scalable technology for measuring genome-wide expression at high spatial resolution. Science. 2019;363(6434):1463–1467.

10. Cleary B, Simonton B, Bezney J, Murray E, Alam S, Sinha A, et al. Compressed sensing for highly efficient imaging transcriptomics. Nat Biotechnol. 2021;39(8):936–942.

11. Zhou X, Seow WY, Ha N, Cheng TH, Jiang L, Boonruangkan J, et al. Highly sensitive spatial transcriptomics using FISHnCHIPs of multiple co-expressed genes. Nat Commun. 2024;15(1):2342.

12. Hemminger Z, Sanchez-Tam G, Ocampo HD, Wang A, Underwood T, Xie F, et al. Spatial single-cell mapping of transcriptional differences across genetic backgrounds in mouse brains. bioRxivorg. 2024;.

13. Yao Z, van Velthoven CTJ, Kunst M, Zhang M, McMillen D, Lee C, et al. A high-resolution transcriptomic and spatial atlas of cell types in the whole mouse brain. Nature. 2023;624(7991):317–332.

14. Qiu C, Martin BK, Welsh IC, Daza RM, L. Tm, Huang X, et al. A single-cell time-lapse of mouse prenatal development from gastrula to birth. Nature. 2024;626(8001):1084–1093.

15. Rinne V, Gröndahl-Yli-Hannuksela K, Fair-Mäkelä R, Salmi M, Rantakari P, Lönnberg T, et al. Single-cell transcriptome analysis of the early immune response in the lymph nodes of Borrelia burgdorferi-infected mice. Microbes and Infection. 2025;27(2):105424.

16. Tibbitt CA, Stark JM, Martens L, Ma J, Mold JE, Deswarte K, et al. Single-cell RNA sequencing of the T helper cell response to house dust mites defines a distinct gene expression signature in airway Th2 cells. Immunity. 2019;51(1):169–184.

17. Sheridan RM, Doan TA, Lucas CJ, Forward TS, Fleming I, Olsen VM, et al. A specific gene expression program underlies antigen archiving by lymphatic endothelial cells in mammalian lymph nodes. Nature Communications. 2025;16(1):8375.

18. Ado S, Dong C, Attaf N, Moussa M, Carrier A, Milpied P, et al. FB5P-seq-mAbs: monoclonal antibody production from FB5P-seq libraries for integrative single-cell analysis of B cells. Frontiers in Immunology. 2024;15:1505971.

19. Hershberg EA, Camplisson CK, Close JL, Attar S, Chern R, Liu Y, et al. PaintSHOP enables the interactive design of transcriptome-and genome-scale oligonucleotide FISH experiments. Nat Methods. 2021;18(8):937–944.

20. Xia C, Babcock HP, Moffitt JR, Zhuang X. Multiplexed detection of RNA using MERFISH and branched DNA amplification. Scientific Reports. 2019;9:7721. doi:10.1038/s41598-019-43943-8.

21. Guan N, Zhang X, Luo Z, Tao D, Yang X. Discriminant projective non-negative matrix factorization. PloS one. 2013;8(12):e83291.

22. Song D, Li K, Hemminger Z, Wollman R, Li JJ. scPNMF: sparse gene encoding of single cells to facilitate gene selection for targeted gene profiling. Bioinformatics. 2021;37(Supplement 1):i358–i366.

23. Raj A, van den Bogaard P, Rifkin SA, van Oudenaarden A, Tyagi S. Imaging individual mRNA molecules using multiple singly labeled probes. Nat Methods. 2008;5(10):877–879.

24. Young AP, Jackson DJ, Wyeth RC. A technical review and guide to RNA fluorescence in situ hybridization. PeerJ. 2020;8:e8806.

25. Yilmaz LS, Noguera DR. Mechanistic approach to the problem of hybridization efficiency in fluorescent in situ hybridization. Appl Environ Microbiol. 2004;70(12):7126–7139.

26. Choi HMT, Schwarzkopf M, Fornace M, Acharya A, Artavanis G, Stegmaier J, et al. Third-generation in situ hybridization chain reaction: multiplexed, quantitative, sensitive, versatile, robust. Development. 2018;145(12):dev165753.

27. Ke R, Mignardi M, Pacureanu A, Svedlund J, Botling J, Wählby C, et al. In situ sequencing for RNA analysis in preserved tissue and cells. Nat Methods. 2013;10(9):857–860.

28. Wang X, Allen WE, Wright MA, Sylwestrak EL, Samusik N, Vesuna S, et al. Three-dimensional intact-tissue sequencing of single-cell transcriptional states. Science. 2018;361(6400).

29. Player AN, Shen LP, Kenny D, Antao VP, Kolberg JA. Single-copy gene detection using branched DNA (bDNA) in situ hybridization. Journal of Histochemistry & Cytochemistry. 2001;49(5):603–611.

30. Biancalani T, Scalia G, Buffoni L, Avasthi R, Lu Z, Sanger A, et al. Deep learning and alignment of spatially resolved single-cell transcriptomes with Tangram. Nat Methods. 2021;18(11):1352–1362.

31. Kleshchevnikov V, Shmatko A, Dann E, Aivazidis A, King HW, Li T, et al. Cell2location maps fine-grained cell types in spatial transcriptomics. Nat Biotechnol. 2022;40(5):661–671.

32. Luecken MD, Büttner M, Chaichoompu K, Danese A, Interlandi M, Mueller MF, et al. Benchmarking atlas-level data integration in single-cell genomics. Nature Methods. 2022;19(1):41–50. doi:10.1038/s41592-021-01336-8.

